# Differential regulation of mitochondrial quality control in skeletal muscle by HZE radiation exposure and partial weightbearing in mice

**DOI:** 10.1101/2025.07.18.665555

**Authors:** Carla MC Nascimento, James D. Fluckey, Florence Lima, Brandon R. Macias, Yasaman Shirazi-Fard, Elizabeth S. Greene, Leslie A. Braby, Susan A. Bloomfield, Michael P Wiggs

## Abstract

Spaceflight places astronauts under both reduced mechanical loading and ionizing radiation, each of which can compromise skeletal muscle health. We investigated whether 21 days of simulated lunar gravity (one sixth G) with or without a single 0.5 Gy dose of ^28^Si heavy ion radiation alters transcriptional regulators of mitochondrial quality control in mouse gastrocnemius muscle. Female BALB/cByJ mice were assigned to four groups: Sham + 1 G (SHAM+CC), Rad + 1 G (RAD+CC), Sham + G/6 (SHAM+G/6), Rad + G/6 (RAD+G/6) and relative mRNA levels of key regulators of mitochondrial biogenesis, mitophagy, dynamics and electron transport chain content were measured by quantitative RT-PCR. Radiation significantly suppressed PGC-1α (p = 0.035) and TFAM (p = 0.051) transcripts and reduced LC3b (p = 0.033) and Park2 (p = 0.007) expression; no effects of simulated lunar gravity or interaction effects were detected. Composite scores confirmed suppression of biogenesis (p = 0.029) and a trend toward reduced mitophagy (p = 0.057). Transcripts encoding oxidative phosphorylation subunits and fusion and fission factors remained unchanged, suggesting preserved mitochondrial content and network homeostasis at day 21. These findings indicate that a single space relevant heavy ion exposure selectively disrupts early transcriptional steps of mitochondrial turnover without immediately altering organelle abundance of transcripts for electron transport chain or dynamics; in contrast simulated lunar gravity alone did not elicit changes in these pathways.

## Introduction

Spaceflight presents a unique physiological challenge to astronauts, as the reduced mechanical loading associated with microgravity and partial-gravity environments can profoundly affect skeletal muscle health. Numerous investigations in humans and rodents have documented decreases in muscle mass, fiber size, and force-generating capacity in response to spaceflight or simulated microgravity (Fitts et al., 2010; LeBlanc et al., 2000). Chronic exposure to reduced loading remains a major risk factor for musculoskeletal deconditioning, and although unloading causes atrophy, it represents only one facet of the musculoskeletal challenges encountered in space.

Astronauts are also exposed to space radiation, particularly high atomic number, high energy (HZE) particles that have high linear energy transfer (LET) from galactic cosmic rays. While skeletal muscle has traditionally been viewed as radioresistant due to its post-mitotic nature, acute doses of ionizing radiation have been shown to induce subtle cellular changes with potential long-term implications (Krause et al., 2017; Shtifman et al., 2013). We previously reported that simulated partial gravity (1/6^th^ G) over 21 days led to significant muscle atrophy, whereas a space-relevant dose of ^28^Si radiation alone did not. Furthermore, there was no synergistic or additive effect of radiation on the atrophy induced by unloading (Wiggs et al., 2023). Despite these null findings at the tissue level, there remains the possibility that radiation exposure could exert more nuanced effects on subcellular processes, particularly those regulating mitochondrial health, without overtly affecting muscle mass.

Mitochondria play a central role in skeletal muscle growth, regeneration, and overall tissue homeostasis (Hyatt & Powers, 2021). Unlike the post-mitotic muscle fiber, mitochondria house their own DNA (mtDNA) and may therefore be more vulnerable to radiation-induced damage (Averbeck & Rodriguez-Lafrasse, 2021). This vulnerability underscores the importance of mitochondrial quality control (MQC), an combination of several processes that include mitochondrial content and turnover (biogenesis and autophagy) and shape/dynamics (fusion and fission) that preserves mitochondrial integrity (Kang et al., 2016; Tilokani et al., 2018). Even in the absence of observable changes in muscle mass, radiation and partial unloading may disrupt MQC pathways, that could potentially have future effects on mitochondrial function and therefore muscle mass and function (Kam & Banati, 2013). Therefore, the present study aimed to investigate whether partial weightbearing (simulating Lunar gravity) with or without ^28^Si radiation alter transcriptional regulators of MQC. We hypothesized that radiation and/or partial loading will disrupt mRNA markers of MQC.

## Materials and Methods

### Animals

All animal procedures were approved by the Institutional Animal Care and Use Committees (IACUC) at both Texas A&M University and Brookhaven National Laboratory (BNL) and have previously been described (Wiggs et al., 2023). Briefly, Four-month-old female BALB/cByJ mice were obtained from Jackson Laboratories (Bar Harbor, Maine). The mice were housed in a controlled environment (23°C ± 2°C, 12:12 hr dark/light cycle) with free access to standard chow and water at BNL. Mice were ranked by body weight and block-assigned to one of two groups: (1) a cage control group (CC), representing normal earthbound gravity (1G), or (2) a partial weight-bearing group simulating lunar gravity (1/6 G). These groups were further divided into subgroups that either received a sham radiation dose or a single whole-body exposure of 0.5 Gy of 300 MeV/u ^28^Si radiation, resulting in four experimental groups: SHAM+CC; RAD+CC; SHAM+G/6; and RAD+/G/6. Radiation exposure took place on Day 0 at NASA’s Space Radiation Laboratory (NSRL) at BNL. Immediately after the sham or real radiation exposure, the mice underwent 21 days of normal weight-bearing or 1/6^th^ weight-bearing.

### Quantitative real-time PCR analysis

Pulverized gastrocnemius muscle (∼10mg) was homogenized using a bead homogenizer, and RNA isolation was performed using TRIzol Reagent (Life Technologies Corp., CA, USA) according to the manufacturer’s instructions. RNA precipitation occurred followed overnight incubation using 50ng/μL of glycogen as co-precipitant to increase the recovery of nucleic acids during precipitation. RNA purity was checked using the Nanodrop Lite Plus Spectrophotometer (ThermoFisher Scientific). RNA was precipitated and resuspended to a concentration of 200ng/μL using RNase-free water. cDNA was synthesized from the RNA samples using a High-Capacity cDNA Reverse Transcription Kit (Applied Biosystems, CA, USA) and was amplified in a 10*μ*L reaction containing appropriate primer pairs and SYBR Green Mastermix as appropriate (Applied Biosystems). SYBR Green primers were designed using Primer-BLAST through PubMed. Custom primer pairs are shown in **Table 1**. The samples were incubated at 95°C for 4 min, followed by 40 cycles of denaturation, annealing, and extension at 95, 60, and 72°C. Fluorescence was measured at the end of the extension step for each cycle using a QuantStudio^TM^ 6 Pro (Applied Biosystems, Waltham, MA). Cycle threshold (Ct) values were determined for each target gene, and expression data were normalized to the geometric mean of two housekeeping genes: β2-microglobulin (β2M) and glyceraldehyde-3-phosphate dehydrogenase (GAPDH). Relative gene expression was calculated using the 2^−ΔCt method and expressed as the normalized fold difference compared to the control group (SHAM +CC). All comparisons were made across experimental conditions involving variations in load and radiation exposure.

**Table 1.**
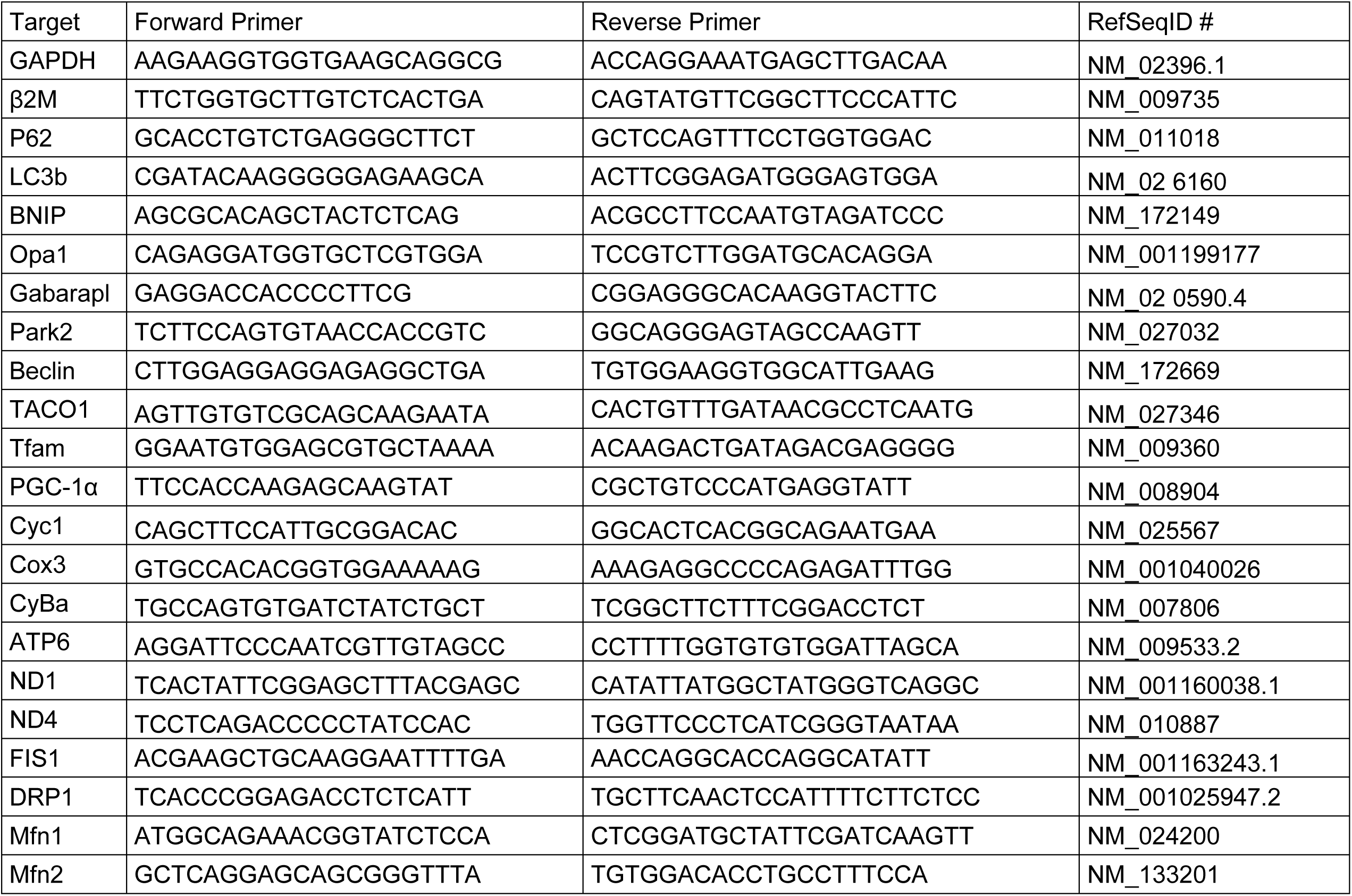
Primer pairs used for SYBR-Green-based quantitative qRT-PCR.

### Statistical analysis

Normalized gene expression values for each of the 21 target transcripts were first assessed for normality (Shapiro–Wilk test) and homogeneity of variance (Levene’s test). If both tests were satisfied (α ≥ 0.05), data were analyzed using Load x Radiation two-way ANOVA. If either test failed, data were power-transformed using Box–Cox (for strictly positive values) or Yeo–Johnson (for zero/negative values).

These procedures automatically estimate γ to maximize normality. Normality and homogeneity of variance were reassessed. Once both criteria were met, the two-way ANOVA was performed. When significant F ratios were found, a Tukey post-hoc test was performed to identify differences within the groups.

To summarize pathway-level effects, genes were grouped into four functional sets (Biogenesis: Nrf2, PGC1a, Tfam, Taco1; Autophagy: Beclin, LC3b, p62, Gabarapl, Park2, Bnip3; Dynamics: Mfn1, Mfn2, Opa1, Fis1, Drp1; OxPhos: COX3, CyBa, CyC1, ATP6, ND4, ND1). For each gene, we first calculated log₂-normalized expression (normalized to the SHAM+CC housekeeping mean). We then z-scored those log₂ values across all samples. Within each pathway, we took the average of the z-scores for every sample to obtain a single composite pathway z-score. These composite scores were then subjected to the same two-way ANOVA as the individual genes.

Data were processed using the following Python libraries: pandas (for data manipulation), NumPy (for numerical operations), SciPy (for statistical tests and transforms), statsmodels (for two-way ANOVA), and scikit-learn (for Yeo– Johnson/Box–Cox transforms). The specific code is available through **supplemental file 1**. SigmaStat v3.4 (Systat Software Inc., San Jose, California) was used for any additional post hoc testing and verification. All figures were compiled using GraphPad Prism 9 (La Jolla, CA). To keep graphs consistent, all data are presented as normalized geometric mean ± standard error of the mean (SEM). Significance was set at α ≤ 0.05, however some results were slightly above this value (e.g. 0.051) and discussed as significant within the text. The results of all statistical analysis and all data that were transformed can be found in **supplemental table 1**.

## Results

We previously reported body weight and muscle size following combined simulated lunar gravity (1/6 G) and heavy ion (HZE) (Wiggs et al., 2023). In general, whole-body radiation exposure of 0.5 Gy of ^28^Si did not independently cause muscle atrophy and did not exacerbate muscle atrophy in partial loading. We conducted a secondary analysis examining gene expression patterns of key MQC regulators, aiming to reveal potential underlying mechanisms that may be overlooked when focusing solely on traditional muscle atrophy outcomes. Due to the use of remaining tissue samples, a complete set of muscles was not available for each animal. As a result, we analyzed a subset of samples per group. Specifically, our current analysis lacks one sample from the SHAM+CC group and two from the RAD+CC group, relative to the original sample size reported previously. Body mass and muscle size for the only animal tissues available for this study can be found in **Table 2**. In summary, partial weight bearing significantly reduced final body mass (both SHAM+G/6 and RAD+G/6 vs. SHAM+CC and RAD+CC; p < 0.001). Gastrocnemius mass showed main effects of load (p < 0.001), radiation (p = 0.03), and their interaction (p < 0.001). However, pairwise tests revealed that G/6 groups had lower muscle mass than CC groups regardless of radiation, and there were no differences between SHAM and RAD within either loading condition. Importantly, the findings related to body weight and muscle mass remain consistent with the outcomes reported in the original study (Wiggs et al., 2023).

**Table 2.** Changes in Body and Muscle Mass in Animals Analyzed for mRNA Expression. Partial weightbearing (*G/6*) resulted in significant body mass loss, whereas normally loaded control animals (CC) maintained their body weight. RAD+G/6 were not different than SHAM+G/6. Gastrocnemius mass analysis revealed significant main effects of loading, radiation, and their interaction. Values are presented as means ± SEM. * Main effect of Load. # Main effect of Radiation, ⁑ Interaction between Radiation and Load. Groups not sharing the same letter are significantly different using a post hoc test (*p* > 0.05).

|  | SHAM+CC<br>(n=8) | RAD+CC<br>(n=7) | SHAM+G/6<br>(n=11) | RAD+G/6<br>(n=10) |
| --- | --- | --- | --- | --- |
| Body Mass on Day 21* | 24.81±1.83<sup>a</sup> | 24.28±1.79<sup>a</sup> | 20.66±1.27<sup>b</sup> | 20.69±1.35<sup>b</sup> |
| % Body Mass Change * | 4.67±0.17<sup>a</sup> | 2.95±1.67<sup>a</sup> | -8.69±3.57<sup>b</sup> | -5.47±1.75<sup>b</sup> |
| Gastrocnemius Mass (mg)*#* | 98.43±2.64<sup>a</sup> | 105.24±3.26<sup>a</sup> | 83.5±1.55<sup>b</sup> | 87.26±2.16<sup>b</sup> |

### ^28^Si Radiation but not simulated lunar gravity alter mRNA markers of mitochondrial turnover

Mitochondrial content is the balance between production of new mitochondria (i.e., biogenesis) and degradation (i.e., mitophagy). A key regulator of mitochondrial biogenesis is peroxisome proliferator-activated receptor gamma coactivator 1-alpha (PGC-1α), which plays a central role in initiating mitochondrial DNA (mtDNA) transcription and replication. PGC-1α exerts its effects by activating nuclear respiratory factors (Nrf1 and Nrf2), which in turn stimulate mitochondrial transcription factor A (Tfam). Tfam is essential for the maintenance of mitochondrial function, as it interacts with critical mitochondrial enzymes to support mtDNA stability and gene expression (Chen et al., 2023). An additional, yet underexplored, aspect of mitochondrial quality control involves the translation of mitochondria-encoded mRNAs. The translational activator of cytochrome c oxidase (Taco1) is essential for the proper assembly and function of cytochrome c oxidase, highlighting its role in maintaining mitochondrial biogenesis (Kremer & Rehling, 2024).

Gene-level analysis revealed a significant main effect of radiation on PGC-1α (p = 0.035) and Tfam (p = 0.051). Pairwise comparisons showed significantly reduced expression of both genes in the HZE + full weightbearing (RAD+CC) group relative to SHAM + full weightbearing, indicating that radiation alone, independent of mechanical load, impairs mRNA markers of mitochondrial biogenesis (Figure 1A).

**Figure 1.**
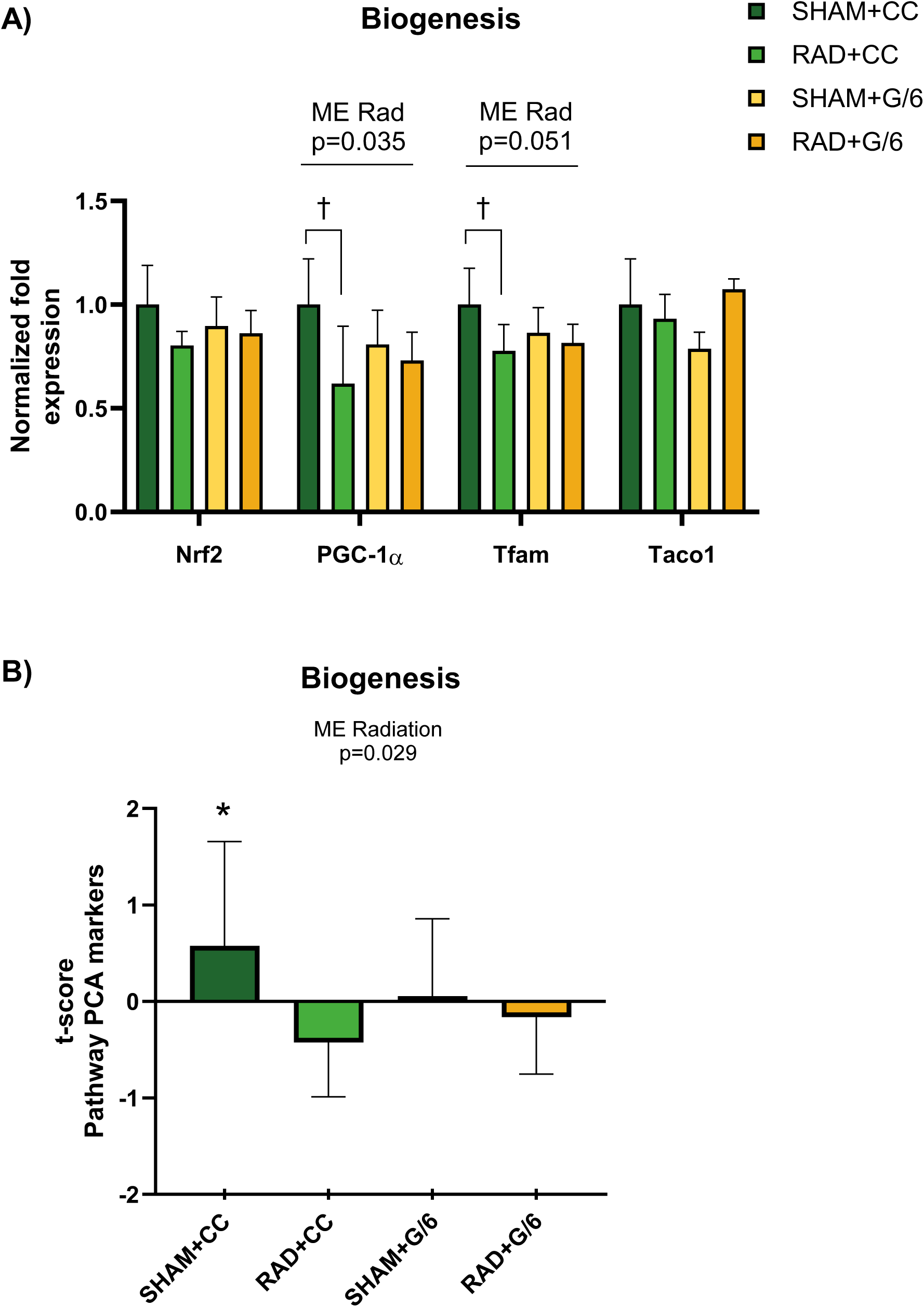
mRNA markers of mitochondrial biogenesis. (A) Relative mRNA levels of key markers mitochondrial biogenesis regulators Nrf2, PGC-1α, Tfam, Taco1 in gastrocnemius muscle from mice under normal loading (CC) or partial weight bearing (G/6), with prior (RAD) ^28^Si radiation or without (SHAM). Two-way ANOVA revealed a significant main effect of radiation on PGC-1α and Tfam, but not on Nrf2 or Taco1. Post hoc pairwise comparisons showed that, under normal loading, irradiated mice (RAD+CC) exhibited lower PGC-1α and Tfam expression compared to sham-irradiated controls (SHAM+CC); denoted by †. No significant differences were observed between G/6 groups. (B) Biogenesis pathway showed a significant main effect of radiation (p=0.029) and pairwise comparisons showed that SHAM+CC was higher than all other groups; denoted by *. Bars represent mean ± SEM.

The composite biogenesis score based on each gene component set (comprising Nrf2, PGC-1α, Tfam, and Taco1) showed a significant main effect of radiation (p=0. 029), indicating lower mitochondrial biogenesis induced by HZE radiation exposure. There was no effect of load (p=0.632) or load × radiation interaction (p=0.152) (Figure 1B).

Mitophagy, a selective form of autophagy, removes damaged mitochondria to preserve cellular homeostasis and regulate mitochondrial content (Onishi et al., 2021). In this study, we examined several key mitophagy-related genes: p62 (also known as SQSTM1), which serves as an adaptor linking ubiquitinated cargo to the autophagosome; LC3b, which drives autophagosome membrane expansion; Bnip3, which targets damaged mitochondria for degradation; and Gabarapl, which supports autophagosome formation and elongation. We also assessed Park2 (Parkin), an E3 ubiquitin ligase that tags defective mitochondria for removal, and Beclin, a critical initiator of autophagosome biogenesis At the individual gene level, LC3b showed a significant main effect of radiation (p = 0.033); pairwise analysis indicated significantly lower expression in the RAD+G/6 group compared to SHAM+G/6 (p = 0.01). Park2 also showed a significant radiation effect (p = 0.007), with decreased expression in both HZE-exposed groups: RAD+CC (p = 0.04) and RAD+G/6 (p = 0.05) relative to their SHAM counterparts. Beclin expression was marginally reduced in RAD+G/6 compared to SHAM+G/6 (p=0.08), suggesting radiation-related suppression under unloading conditions. BNIP3 displayed a trend toward a radiation × load interaction (p = 0.052), hinting at context-specific modulation that warrants further exploration (Figure 2A).

**Figure 2.**
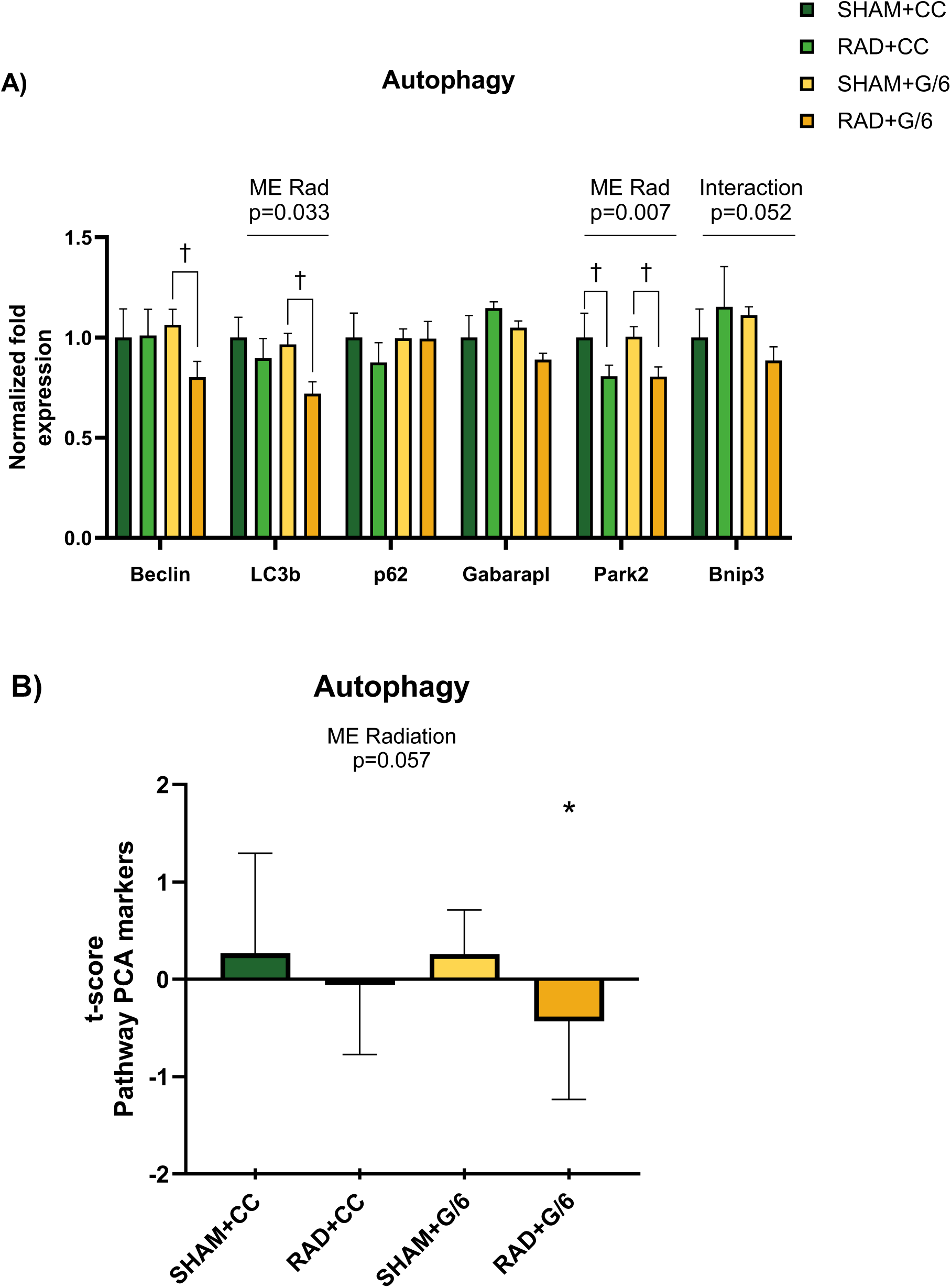
mRNA markers of autophagy. (A) Relative mRNA levels of key markers mitochondrial autophagy regulators Beclin, LC3b, P62, Gabarapl, Park2 and Bnip3 in gastrocnemius muscle from mice under normal loading (CC) or partial weight bearing (G/6), with (RAD) or without (SHAM) ^28^Si radiation. Two-way ANOVA revealed a significant main effect of radiation on LC3b and Park2, and interaction effect on Bnip3, but not on p62 or Gabarapl. Post hoc pairwise comparisons showed that, under partial loading (G/6), irradiated mice (RAD+G/6) exhibited higher Beclin and LC3b expression compared to sham-irradiated controls (SHAM+G/6; denoted by † between comparisons). No significant differences were observed between CC groups. Regarding Park2 expression, both irradiated groups (RAD+CC and RAD+G/6) presented lower gene expression. (B) Autophagy pathway showed a borderline-significant main effect of radiation (p=0.057). where SHAM+G/6 is less than all other groups; denoted by *. Bars represent mean ± SEM.

For the autophagy-pathway composite, a borderline main effect (p=0.057) indicated that radiation trend to reduce the expression of this pathway. There were no effects of load (p=0.464) or the interaction term (p=0.486) (Figure 2B).

Although mitochondrial biogenesis and autophagy gene expression inform us about turnover rates, they do not directly quantify the existing mitochondrial pool. To estimate total mitochondrial content, we therefore measured mRNA levels of oxidative phosphorylation (OXPHOS) complex subunits encoded by the mitochondrial genome. In skeletal muscle, steady-state abundance of these transcripts often correlates with mitochondrial volume and density (Malik & Czajka, 2013; Short et al., 2005), since each mitochondrion carries its own complement of mtDNA-encoded OXPHOS genes .The selected genes represent components of multiple electron transport chain (ETC) complexes and include both mitochondrial DNA (mtDNA)-encoded and nuclear DNA (nDNA)-encoded transcripts. Specifically, we evaluated: Complex I: ND1 and ND4 (mtDNA-encoded); Complex III: Cytochrome b (CyBa, mtDNA-encoded) and Cyc1 (nDNA-encoded), a subunit implicated in electron transfer and ROS production (Bleier & Drose, 2013); Complex IV: COX3 (mtDNA-encoded); Complex V: ATP6 (mtDNA-encoded), involved in ATP synthesis. Neither the pathway-level analysis (Figure 3A) nor the individual gene-level analysis (Figure 3B) revealed significant differences across experimental groups. No significant changes were observed in any of the OXPHOS transcripts across loading or radiation conditions, suggesting that neither intervention altered total mitochondrial content as estimated at the mRNA level.

**Figure 3.**
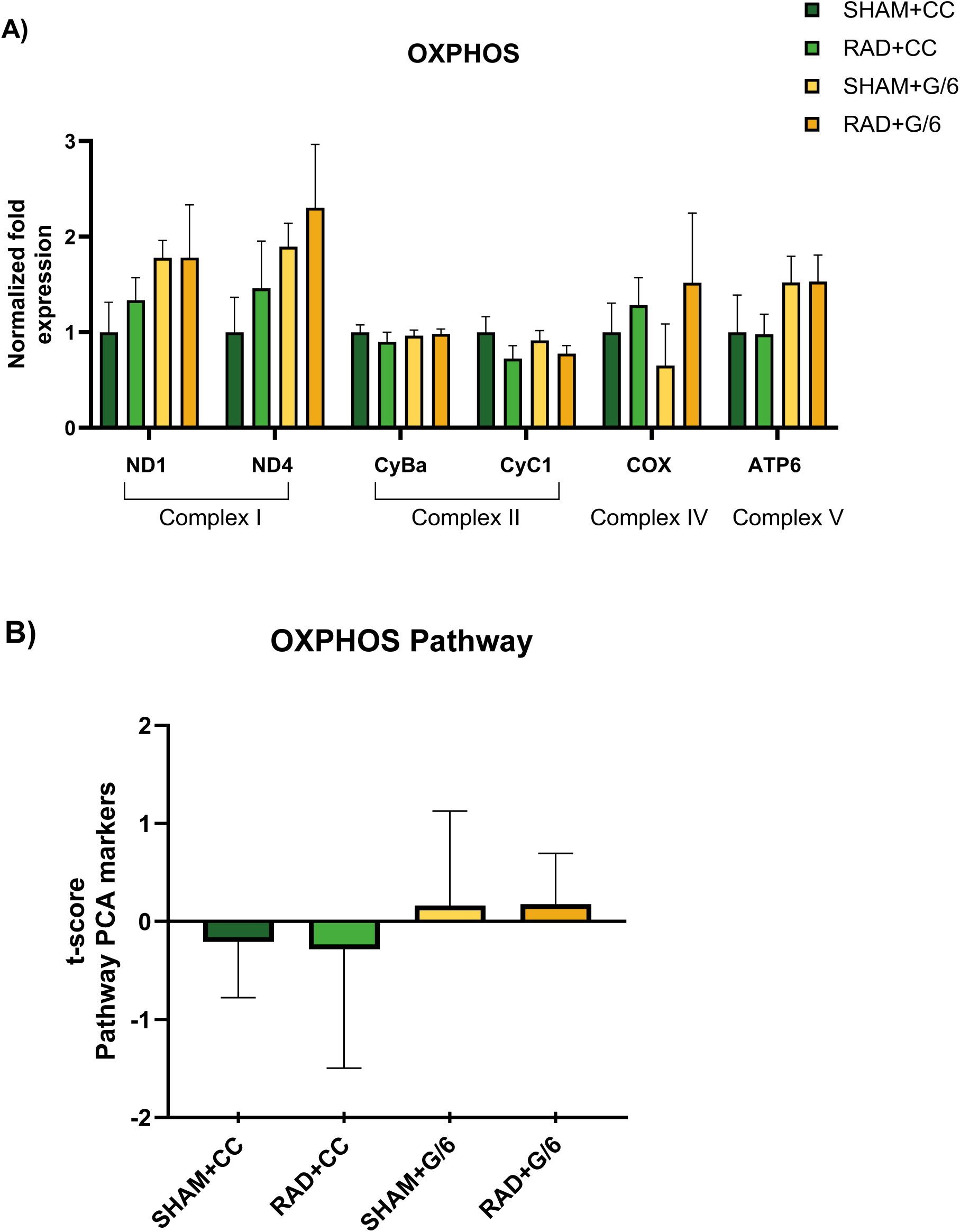
mRNA markers of mitochondrial OXPHOS. These genes are among the 13 protein-coding genes in mitochondrial DNA (mtDNA). (A) Relative mRNA levels of key markers mitochondrial OXPHOS regulators for Complex I: ND-1, ND-4; Complex III: CyBa, CyC; Complex IV: COX3, and Complex V: ATP-6 in gastrocnemius muscle from mice under normal loading (CC) or partial weight bearing (G/6), with (RAD) or without (SHAM) ^28^Si radiation. Bars represent mean ± SEM. No significant main effects or pairwise differences were observed between groups. (B) OXPHOS pathway also showed no significant differences.

The final component of mitochondrial quality control is dynamics, which encompass the opposing processes of fusion and fission that remodel the mitochondrial network. Fusion, mediated by mitofusins (Mfn1/2) and Opa1, allows healthy mitochondria to mix contents and dilute damage, whereas fission, driven by Drp1 and Fis1, facilitates segregation and removal of dysfunctional segments; an imbalance between these processes may lead to the accumulation of defective organelles. To assess potential changes in this system, we conducted a pathway-level analysis of fusion and fission gene expression. Our results revealed no significant differences across experimental groups (Figure 4A). Mitochondrial fusion is regulated by Mitofusin 1 and 2 (Mfn1, Mfn2) in the outer membrane and Optic atrophy 1 (Opa1) in the inner membrane, all of which are critical for mitochondrial remodeling and function (Tilokani et al., 2018). However, we observed no significant changes in Mfn1, Mfn2, or Opa1 expression among the groups (Figure 4B). Similarly, mitochondrial fission—controlled by Dynamin-related protein 1 (Drp1) and Fission protein 1 (Fis1)—plays a key role in mitochondrial turnover and mitophagy (Liu et al., 2024). Despite evidence linking mitofusin deficiency to reduced mtDNA copy number and mitochondrial myopathy (Chen et al., 2010), our data showed no significant alterations in Drp1 or Fis1 expression (Figure 4B). Together, these findings suggest that mitochondrial fusion and fission mechanisms remain stable under the tested conditions, with no significant changes in the expression of key regulatory genes.

**Figure 4.**
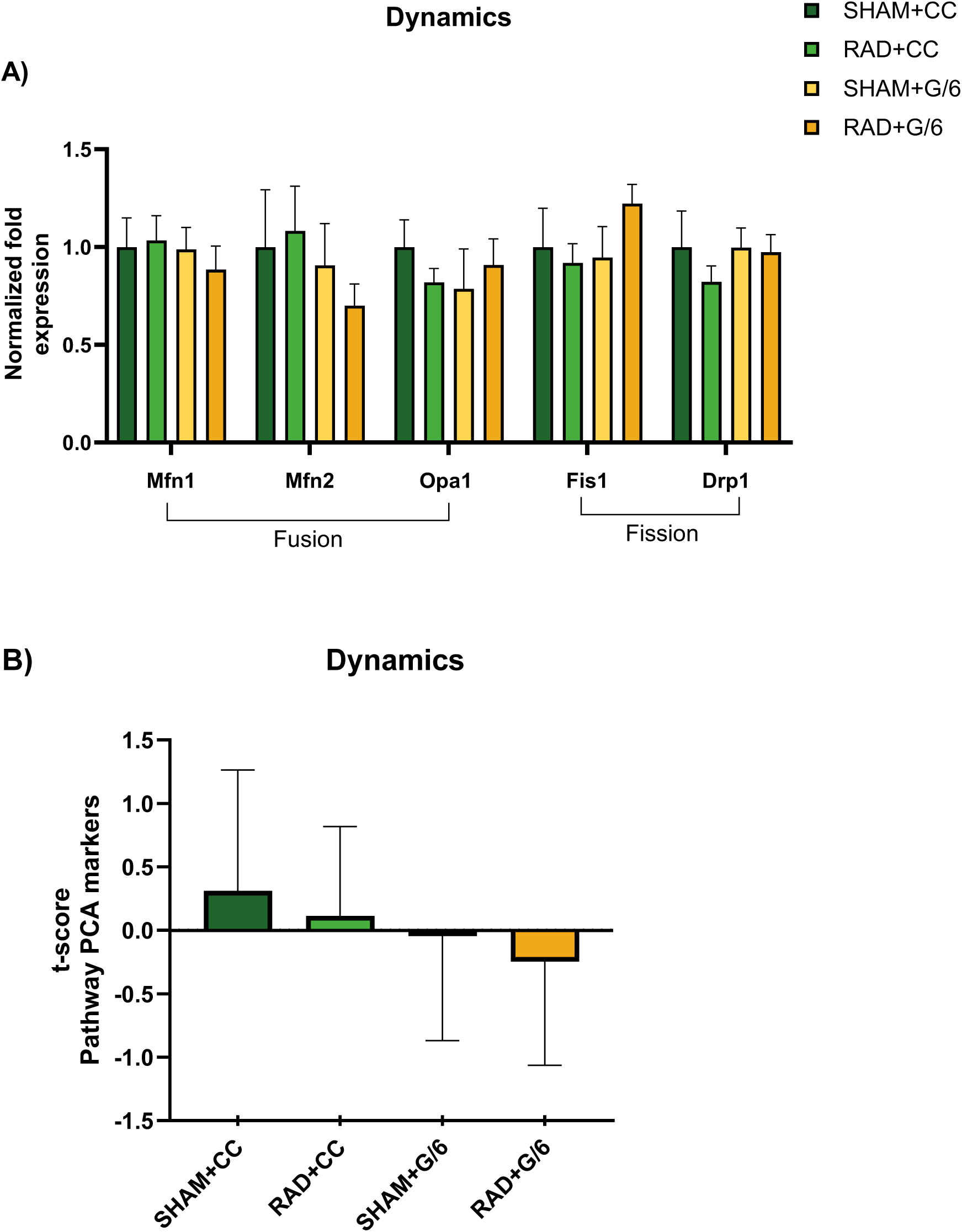
mRNA markers of mitochondrial Dynamics – Fusion and Fission. (A) Relative mRNA levels of key markers mitochondrial Fusion regulators Mfn1, Mfn2 and Opa1 and Fission Fis1 and Drp1 in gastrocnemius muscle from mice under normal loading (CC) or partial weight bearing (G/6), with (RAD) or without (SHAM) ^28^Si radiation. No significant main effects or pairwise differences were observed between groups. (B) Mitochondrial Dynamics pathway also showed no significant differences. Bars represent mean ± SEM.

## Discussion

In this study, we investigated whether a single dose of 0.5 Gy of 300 MeV/u ^28^Si heavy-ion radiation with or without simulated lunar gravity (partial weight bearing, G/6) and would alter the transcriptional landscape of mitochondrial quality control (MQC) pathways in murine skeletal muscle. Our primary observations were that radiation, but not partial unloading, suppressed the mRNA markers of both mitochondrial biogenesis (PGC-1α, Tfam) and mitophagy (LC3b, Park2) on day 21. In contrast, neither intervention elicited measurable changes in transcript-level proxies of mitochondrial content (OXPHOS subunits) or in the expression of genes governing mitochondrial fusion and fission. These findings suggest that acute heavy-ion radiation may selectively disrupt transcriptional steps associated with mitochondrial turnover, while partial weightbearing under lunar gravity conditions may not elicit a sustained gene-level response at this time point.

Our finding that ^28^Si radiation, similar to predicted exposure for Mars Missions (Cucinotta & Durante, 2006), reduced biogenesis and mitophagy transcripts in the absence of overt muscle atrophy adds to a growing but complex body of literature investigating radiation effects on skeletal muscle. Although our observed downregulation of PGC-1α and Tfam suggests suppression of mitochondrial biogenesis, prior studies have reported variable outcomes depending on radiation type, dose, timing, and tissue examined. For example, 24 hours post-irradiation, exposure to 2 Gy of 260 kvp x rays increased markers of mitochondrial biogenesis and content in muscle, consistent with a transient oxidative stress response (Kim et al., 2019). In contrast, Hardee et al. found no changes in PGC-1α, Tfam, or LC3b two weeks following localized exposure to 16 Gy of 320kvp x rays, suggesting a potential dose threshold above which skeletal muscle either fails to mount or bypasses a transcriptional response (Hardee et al., 2014). When hindlimb unloading was combined with gamma irradiation, reductions were observed in genes such as ND1, ND4, COX3, COX8B, and ATP5G1, although the absence of a radiation-only group makes it difficult to isolate the specific effects of irradiation (Tran & Choi, 2022).

Compared to the broader literature on low-LET or high-dose exposures, relatively little is known about how low-dose, high-LET radiation in the form of HZE particles encountered in space impacts mitochondrial pathways in post-mitotic tissue. This represents a critical gap in our understanding, as skeletal muscle may lack the proliferative capacity needed for canonical stress responses seen in other cell types. Notably, long-term mitochondrial dysregulation has been reported months after whole-body irradiation in rhesus macaques, even in the absence of marked atrophy, suggesting that radiation-induced changes in mitochondrial turnover may persist over time and carry functional consequences (Lee et al., 2025). These findings support the possibility that space-relevant radiation may initiate early suppression of mitochondrial quality control gene networks that, if sustained, could compromise muscle health under prolonged exposure scenarios (Rudolf & Hood, 2024).

Comparison of our partial weightbearing model with complete hindlimb unloading reveals important differences in the regulation of mitochondrial quality control. In rodent hindlimb suspension studies, transcriptional activation of mitophagy and suppression of biogenesis are evident within the first one to two weeks and the majority of these markers persist (Rosa-Caldwell et al., 2020) (Baehr et al., 2017; Sacheck et al., 2007; Wagatsuma et al., 2011). By contrast, our G/6 model showed no persistent changes in these transcriptional pathways on day 21, suggesting that partial gravity provides sufficient mechanical stimulus to avoid the acute MQC perturbations observed under total unloading. These findings support the notion that the severity and duration of mechanical unloading dictate the temporal dynamics of MQC gene expression, with partial loading producing a milder phenotype that may resolve before our measurement window.

Although radiation suppressed transcripts associated with biogenesis and mitophagy, mRNA levels of oxidative phosphorylation subunits remained unchanged across all groups. This apparent disconnect may arise because reduced biogenesis and autophagy offset one another, maintaining steady mitochondrial content; moreover, recent data in other tissues demonstrate that many mitochondrial proteins, particularly those in the electron transport chain, have unusually long half-lives (Bomba-Warczak & Savas, 2022; Krishna et al., 2021). Transcripts for fusion and fission factors also remained unchanged, indicating that neither ^28^Si radiation nor partial unloading triggered changes in transcriptional markers of mitochondrial dynamics. It remains unclear whether mitochondrial dynamics are altered under these conditions; if fusion or fission are affected, those changes do not appear to involve altered mRNA levels but may instead occur through post-translational modifications such as phosphorylation, acetylation or ubiquitination (Sabouny & Shutt, 2020).

Several limitations of this study should be acknowledged. First, we measured only mRNA levels of markers that are commonly associated with MQC. Without corresponding protein, enzymatic activity or morphological assessments, we cannot confirm whether transcriptional changes translated into functional deficits. Second, sampling at a single 21-day time point may have missed earlier or later transcriptional fluctuations; transient perturbations in MQC pathways are well documented during the first one to two weeks of unloading (Baehr et al., 2017; Sacheck et al., 2007) and may likewise occur after radiation exposure. Finally, our use of an acute 0.5 Gy radiation dose does not fully recapitulate the chronic, fractionated exposure encountered during long-duration spaceflight. Mixed-ion, low-dose-rate models would provide a more accurate simulation of deep-space conditions and may yield different patterns of gene expression.

Our limited sample availability and single time point design prevent evaluation of acute responses to the 0.5 Gy 300 MeV/u 28Si exposure and do not allow functional assays that might explain why mitochondrial gene regulation remained altered 21 days post exposure. Because we did not observe changes in mtDNA encoded genes and the downregulated MQC genes are nuclear encoded across multiple chromosomes, it is very unlikely that HZE tracks at this dosage caused the coordinated DNA damage required to suppress their expression. It is much more likely that a nontargeted mechanism such as an redox process or relatively stable radiation chemistry product that can migrate to mitochondria and damage their membranes is involved (Li et al., 2014). This speculation is consistent with recent evidence implicating membrane targets in persistent space radiation effects in other tissues (Straume et al., 2025).

## Conclusion

In conclusion, our findings suggest that space-relevant radiation can suppress transcriptional markers of mitochondrial biogenesis and mitophagy in skeletal muscle without inducing immediate changes in mitochondrial content or dynamics. Simulated lunar gravity, in contrast, does not elicit sustained transcriptional changes in these pathways on day 21, although transient early effects cannot be ruled out.

These results support the hypothesis that even a single radiation exposure may alter the muscle’s capacity for mitochondrial turnover, which over time could contribute to cumulative organelle dysfunction. Future work should prioritize integrative assessments of mitochondrial quality using protein-level, enzymatic, and morphological metrics, coupled with longitudinal sampling and countermeasure testing to preserve muscle health during extended space missions.

## List of Abbreviations

1/6 G – One-sixth Earth gravity 1G – Normal Earth gravity

2^−ΔCt - Fold-difference expression measure calculated by the 2^−ΔCt method

28Si – Silicon-28 ion

ANOVA – Analysis of variance ATP6 – ATP synthase Fo subunit 6

BNL – Brookhaven National Laboratory BNIP3 – BCL2 Interacting Protein 3 CC – Cage control

COX3 – Cytochrome c oxidase subunit 3 Ct – Cycle threshold

Cytc1 – Cytochrome c1

Drp1 – Dynamin-related protein 1 ETC – Electron transport chain Fis1 – Fission protein 1

G/6 - Partial weightbearing simulating lunar gravity (one sixth G) Gabarapl – Gamma-aminobutyric acid receptor-associated protein-like GAPDH – Glyceraldehyde-3-phosphate dehydrogenase

HZE – High charge (Z) and high energy (heavy ion radiation) IACUC – Institutional Animal Care and Use Committees

LET – Linear energy transfer

LC3b - Microtubule-associated proteins 1A/1B light chain 3 (isoform b) ME – main effect

Mfn1 – Mitofusin 1 Mfn2 – Mitofusin 2

MQC – Mitochondrial quality control mtDNA – Mitochondrial DNA

NASA – National Aeronautics and Space Administration

ND1 - NADHSub1 - NADH dehydrogenase subunit 1 (Complex I) ND4 - NADHSub4 - NADH dehydrogenase subunit 4 (Complex I) NRF2 - Nuclear respiratory factor 2

NSRL - NASA’s Space Radiation Laboratory Opa1 – Optic atrophy 1

OXPHOS – Oxidative phosphorylation qRT-PCR – quantitative real-time PCR P62 – Sequestosome 1 (SQSTM1)

Park2 – Parkin RBR E3 ubiquitin protein ligase

PGC-1α – Peroxisome proliferator-activated receptor gamma coactivator 1-alpha RAD – groups exposed to ^28^Si radiation

ROS – Reactive oxygen species SEM – Standard error of the mean

SHAM – Sham-irradiated (control radiation) group

TACO1 – Translational activator of cytochrome c oxidase subunit I Tfam – Mitochondrial transcription factor A

## Acknowledgements

The authors would like to thank all members of the Muscle Biology Lab (PI: JDF), Bone Biology Lab (PI: SAB) for their support and assistance. We are grateful for the expert assistance of the BNL Medical Department staff (M. Petry, K. Bonti, L. Thompson and P. Guida) and NSRL physicists (A. Rusek and M. Sievertz), which was essential to the success of our NSRL experiments at BNL. These studies were funded through the NASA Cooperative Agreement NCC 9-58 with the National Space Biomedical Research Institute (no. NCC 9-58-MA01602 to SAB). Support was also provided by a National Space Biomedical Research Institute Graduate Training Fellowship (no. NSBRI-RFP-05-02 to BRM)

## Authors Contributions

**CMCN** – conceptualization, formal analysis, investigation, validation, visualization, writing- original draft preparation. **JDF** – conceptualization, data curation, funding acquisition, investigation, resources, writing – review and editing. **FL** – conceptualization, data curation, investigation, methodology, writing- review and editing, **BRM** - conceptualization, data curation, investigation, writing- review and editing, **YS** - conceptualization, data curation, investigation, writing- review and editing; **ESG** - investigation, methodology, writing- review and editing, **LAB** – conceptualization, funding acquisition, supervision, writing – review and editing, **SAB** - conceptualization, data curation, funding acquisition, investigation, methodology, project administration, resources supervision, writing – review and editing. **MPW** - conceptualization, data curation, formal analysis, investigation, resources, software, validation, visualization, writing- original draft preparation.

## References

1. Averbeck, D., & Rodriguez-Lafrasse, C. (2021). Role of Mitochondria in Radiation Responses: Epigenetic, Metabolic, and Signaling Impacts. Int J Mol Sci, 22(20). 10.3390/ijms222011047

2. Baehr, L. M., West, D. W., Marshall, A. G., Marcotte, G. R., Baar, K., & Bodine, S. C. (2017). Muscle-specific and age-related changes in protein synthesis and protein degradation in response to hindlimb unloading in rats. Journal of Applied Physiology, 122(5), 1336–1350.

3. Bleier, L., & Drose, S. (2013). Superoxide generation by complex III: from mechanistic rationales to functional consequences. Biochim Biophys Acta, 1827(11-12), 1320–1331. 10.1016/j.bbabio.2012.12.002

4. Bomba-Warczak, E., & Savas, J. N. (2022). Long-lived mitochondrial proteins and why they exist. Trends in cell biology, 32(8), 646–654.

5. Chen, H., Vermulst, M., Wang, Y. E., Chomyn, A., Prolla, T. A., McCaffery, J. M., & Chan, D. C. (2010). Mitochondrial fusion is required for mtDNA stability in skeletal muscle and tolerance of mtDNA mutations. Cell, 141(2), 280–289. 10.1016/j.cell.2010.02.026

6. Chen, X., Ji, Y., Liu, R., Zhu, X., Wang, K., Yang, X., Liu, B., Gao, Z., Huang, Y., Shen, Y., Liu, H., & Sun, H. (2023). Mitochondrial dysfunction: roles in skeletal muscle atrophy. J Transl Med, 21(1), 503. 10.1186/s12967-023-04369-z

7. Cucinotta, F. A., & Durante, M. (2006). Cancer risk from exposure to galactic cosmic rays: implications for space exploration by human beings. The lancet oncology, 7(5), 431–435.

8. Fitts, R. H., Trappe, S. W., Costill, D. L., Gallagher, P. M., Creer, A. C., Colloton, P. A., Peters, J. R., Romatowski, J. G., Bain, J. L., & Riley, D. A. (2010). Prolonged space flight-induced alterations in the structure and function of human skeletal muscle fibres. J Physiol, 588(Pt 18), 3567–3592. 10.1113/jphysiol.2010.188508

9. Hardee, J. P., Puppa, M. J., Fix, D. K., Gao, S., Hetzler, K. L., Bateman, T. A., & Carson, J. A. (2014). The effect of radiation dose on mouse skeletal muscle remodeling. Radiology and oncology, 48(3), 247.

10. Hyatt, H. W., & Powers, S. K. (2021). Mitochondrial Dysfunction Is a Common Denominator Linking Skeletal Muscle Wasting Due to Disease, Aging, and Prolonged Inactivity. Antioxidants (Basel), 10(4). 10.3390/antiox10040588

11. Kam, W. W.-Y., & Banati, R. B. (2013). Effects of ionizing radiation on mitochondria. Free Radical Biology and Medicine, 65, 607–619.

12. Kang, C., Yeo, D., & Ji, L. L. (2016). Muscle immobilization activates mitophagy and disrupts mitochondrial dynamics in mice. Acta Physiol (Oxf), 218(3), 188–197. 10.1111/apha.12690

13. Kim, E. J., Lee, M., Kim, D. Y., Kim, K. I., & Yi, J. Y. (2019). Mechanisms of energy metabolism in skeletal muscle mitochondria following radiation exposure. Cells, 8(9), 950.

14. Krause, A. R., Speacht, T. L., Zhang, Y., Lang, C. H., & Donahue, H. J. (2017). Simulated space radiation sensitizes bone but not muscle to the catabolic effects of mechanical unloading. PLoS One, 12(8), e0182403. 10.1371/journal.pone.0182403

15. Kremer, L. S., & Rehling, P. (2024). Coordinating mitochondrial translation with assembly of the OXPHOS complexes. Hum Mol Genet, 33(R1), R47–R52. 10.1093/hmg/ddae025

16. Krishna, S., Drigo, R. A., Capitanio, J. S., Ramachandra, R., Ellisman, M., & Hetzer, M. W. (2021). Identification of long-lived proteins in the mitochondria reveals increased stability of the electron transport chain. Developmental cell, 56(21), 2952–2965. e2959.

17. LeBlanc, A., Schneider, V., Shackelford, L., West, S., Oganov, V., Bakulin, A., & Voronin, L. (2000). Bone mineral and lean tissue loss after long duration space flight. J Musculoskelet Neuronal Interact, 1(2), 157–160. https://www.ncbi.nlm.nih.gov/pubmed/15758512

18. Lee, J., Chen, X., Fanning, K. M., Si, C., Davis, A. T., Wasserman, D. H., Bracy, D., Furdui, C. M., & Kavanagh, K. (2025). Persistent Postirradiation Skeletal Muscle Protein and Insulin Sensitivity Changes in Nonhuman Primates. Radiation research.

19. Li, M., Gonon, G., Buonanno, M., Autsavapromporn, N., De Toledo, S. M., Pain, D., & Azzam, E. I. (2014). Health risks of space exploration: targeted and nontargeted oxidative injury by high-charge and high-energy particles. Antioxidants & redox signaling, 20(9), 1501–1523.

20. Liu, B. H., Xu, C. Z., Liu, Y., Lu, Z. L., Fu, T. L., Li, G. R., Deng, Y., Luo, G. Q., Ding, S., Li, N., & Geng, Q. (2024). Mitochondrial quality control in human health and disease. Mil Med Res, 11(1), 32. 10.1186/s40779-024-00536-5

21. Malik, A. N., & Czajka, A. (2013). Is mitochondrial DNA content a potential biomarker of mitochondrial dysfunction? Mitochondrion, 13(5), 481–492.

22. Onishi, M., Yamano, K., Sato, M., Matsuda, N., & Okamoto, K. (2021). Molecular mechanisms and physiological functions of mitophagy. EMBO J, 40(3), e104705. 10.15252/embj.2020104705

23. Rosa-Caldwell, M. E., Brown, J. L., Perry Jr, R. A., Shimkus, K. L., Shirazi-Fard, Y., Brown, L. A., Hogan, H. A., Fluckey, J. D., Washington, T. A., & Wiggs, M. P. (2020). Regulation of mitochondrial quality following repeated bouts of hindlimb unloading. Applied Physiology, Nutrition, and Metabolism, 45(3), 264–274.

24. Rudolf, A. M., & Hood, W. R. (2024). Mitochondrial stress in the spaceflight environment. Mitochondrion, 101855.

25. Sabouny, R., & Shutt, T. E. (2020). Reciprocal regulation of mitochondrial fission and fusion. Trends in biochemical sciences, 45(7), 564–577.

26. Sacheck, J. M., Hyatt, J. P. K., Raffaello, A., Thomas Jagoe, R., Roy, R. R., Reggie Edgerton, V., Lecker, S. H., & Goldberg, A. L. (2007). Rapid disuse and denervation atrophy involve transcriptional changes similar to those of muscle wasting during systemic diseases. The FASEB Journal, 21(1), 140–155.

27. Short, K. R., Bigelow, M. L., Kahl, J., Singh, R., Coenen-Schimke, J., Raghavakaimal, S., & Nair, K. S. (2005). Decline in skeletal muscle mitochondrial function with aging in humans. Proceedings of the National Academy of Sciences, 102(15), 5618–5623.

28. Shtifman, A., Pezone, M. J., Sasi, S. P., Agarwal, A., Gee, H., Song, J., Perepletchikov, A., Yan, X., Kishore, R., & Goukassian, D. A. (2013). Divergent modification of low-dose (5)(6)Fe-particle and proton radiation on skeletal muscle. Radiat Res, 180(5), 455–464. 10.1667/RR3329.1

29. Straume, T., Mora, A. M., Brown, J. B., Bansal, I., Rabin, B. M., Braby, L. A., & Wyrobek, A. J. (2025). Non-DNA radiosensitive targets that initiate persistent behavioral deficits in rats exposed to space radiation. Life Sciences in Space Research, 45(C).

30. Tilokani, L., Nagashima, S., Paupe, V., & Prudent, J. (2018). Mitochondrial dynamics: overview of molecular mechanisms. Essays Biochem, 62(3), 341–360. 10.1042/EBC20170104

31. Tran, K. N., & Choi, J.-i. (2022). Mimic microgravity effect on muscle transcriptome under ionizing radiation. Life Sciences in Space Research, 32, 96–104.

32. Wagatsuma, A., Kotake, N., Kawachi, T., Shiozuka, M., Yamada, S., & Matsuda, R. (2011). Mitochondrial adaptations in skeletal muscle to hindlimb unloading. Molecular and Cellular Biochemistry, 350, 1–11.

33. Wiggs, M. P., Lee, Y., Shimkus, K. L., O’Reilly, C. I., Lima, F., Macias, B. R., Shirazi-Fard, Y., Greene, E. S., Hord, J. M., Braby, L. A., Carroll, C. C., Lawler, J. M., Bloomfield, S. A., & Fluckey, J. D. (2023). Combined effects of heavy ion exposure and simulated Lunar gravity on skeletal muscle. Life Sci Space Res (Amst), 37, 39–49. 10.1016/j.lssr.2023.02.003

34. Wildman, R. P., Muntner, P., Reynolds, K., McGinn, A. P., Rajpathak, S., Wylie-Rosett, J., & Sowers, M. R. (2008). The obese without cardiometabolic risk factor clustering and the normal weight with cardiometabolic risk factor clustering: prevalence and correlates of 2 phenotypes among the US population (NHANES 1999-2004). Archives of internal medicine, 168(15), 1617–1624.

