## Supplemental File 1 for "Differential regulation of mitochondrial quality control in skeletal muscle by HZE radiation exposure and partial weightbearing in mice"

**Supplemental File 1. Python code for data transformation and analysis.**

```
#!/usr/bin/env python3
# qpcr_powertransform_anova.py

import pandas as pd
import numpy as np
from scipy.stats import gmean, shapiro, levene, boxcox, boxcox_normmax
from sklearn.preprocessing import PowerTransformer
from scipy.stats import zscore
from sklearn.decomposition import PCA
import statsmodels.formula.api as smf
from statsmodels.stats.anova import anova_lm
from statsmodels.stats.multicomp import pairwise_tukeyhsd

# 1) Load raw CT data
INPUT_CSV = "plates_rename.csv"
df = pd.read_csv(INPUT_CSV)

# 2) Compute control-group (CC+Sham) mean Cq for each gene
ctrl = df[(df['load']=="CC") & (df['radiation']=="SHAM")]
genes = ['GAPDH','b-ACT',
        'NRF-2','PCG1-a','TFAM',
        'Beclin','LC3b','P62','Gabarapl','Park2','BNIP',
        'Mfn1','Mfn2','Opa1','FIS-1','DRP-1',
        'TACO-1','COX','CyBa','CyC1','ATP6','NADHSub4','NADHsub1']
for g in genes:
    df[f"Cq_ctrl_{g}"] = ctrl[g].mean()

# 3) ΔCq and RQ for each gene
for g in genes:
    df[f"dCt_{g}"] = df[f"Cq_ctrl_{g}"] - df[g]
    df[f"RQ_{g}"] = 2 ** (-df[f"dCt_{g}"])

# 4) Norm factor = geometric mean of housekeeping RQs
df['NormFactor'] = gmean(df[['RQ_GAPDH','RQ_b-ACT']], axis=1)

# 5) Normalized RQ and log2FC for targets
targets = [g for g in genes if g not in ('GAPDH','b-ACT')]
for g in targets:
    df[f"RQn_{g}"] = df[f"RQ_{g}"] / df['NormFactor']
    safe = df[f"RQn_{g}"].replace({0: np.nan})
    df[f"log2FC_{g}"] = np.log2(safe)

# 6) Ensure categorical factors
df['load']      = df['load'].astype('category')
df['radiation'] = df['radiation'].astype('category')

# 7) Gene-by-gene analysis with power transforms, ANOVA, Tukey HSD
print("\n=== GENE-LEVEL ANALYSIS ===")
```

```
for g in targets:
    col = f"log2FC_{g}"
    sub = df.dropna(subset=[col,'load','radiation']).copy()
    if sub.shape[0] < 4:
        print(f"\nSkipping {g}: too few samples")
        continue

    # 7a) Test normality & homogeneity
    sw_p = shapiro(sub[col])[1]
    lev_p = levene(
        *[sub.loc[sub.load==lv, col]
          for lv in sub.load.unique()],
        center='median'
    ).pvalue

    use_col = col
    if sw_p < 0.05 or lev_p < 0.05:
        # need power transform
        x = sub[col].values
        if np.all(x > 0):
            # Box–Cox
            lam = boxcox_normmax(x, method='mle')
            sub[f"{col}_t"] = boxcox(x, lam)
            use_col = f"{col}_t"
            print(f"\n{g}: Box–Cox λ={lam:.2f}")
        else:
            # Yeo–Johnson
            pt = PowerTransformer(method='yeo-johnson', standardize=False)
            sub[f"{col}_t"] = pt.fit_transform(x.reshape(-1,1)).flatten()
            use_col = f"{col}_t"
            print(f"\n{g}: Yeo–Johnson λ={pt.lambdas_[0]:.2f}")

        # re-test (optional)
        sw_p2 = shapiro(sub[use_col])[1]
        lev_p2 = levene(
            *[sub.loc[sub.load==lv, use_col]
              for lv in sub.load.unique()],
            center='median'
        ).pvalue
        print(f"  post-transform Shapiro p={sw_p2:.3f}, Levene p={lev_p2:.3f}")
    else:
        print(f"\n{g}: normal (Shapiro p={sw_p:.3f}, Levene p={lev_p:.3f})")

    # 7b) Group means ± SEM
    stats = sub.groupby(['load','radiation'])[use_col]\
              .agg(mean='mean', sem='sem')\
              .reset_index()
    print(f"\n-- {g} group stats --\n{stats.to_string(index=False)}")

    # 7c) Two-way ANOVA (Type III)
```

```
    model = smf.ols(f"{use_col} ~ C(load)*C(radiation)", data=sub).fit()
    aov = anova_lm(model, typ=3)
    print(f"\n-- {g} ANOVA --\n{aov[['sum_sq','df','F','PR(>F)']]}")

    # 7d) Tukey HSD if any main or interaction p<0.05
    if (aov.loc['C(load)','PR(>F)'] < 0.05
        or aov.loc['C(radiation)','PR(>F)'] < 0.05
        or aov.loc['C(load):C(radiation)','PR(>F)'] < 0.05):
        mc = pairwise_tukeyhsd(sub[use_col],
                              sub['load'].astype(str) + "_" + sub['radiation'])
        print(f"\n-- {g} Tukey HSD --\n{mc.summary()}")

# 8) Pathway-level Z-score composites and ANOVA
groups = {
    'Biogenesis': ['log2FC_NRF-2','log2FC_PCG1-a','log2FC_TFAM','log2FC_TACO-1'],
    'Autophagy' : ['log2FC_Beclin','log2FC_LC3b','log2FC_P62',
                   'log2FC_Gabarapl','log2FC_Park2','log2FC_BNIP'],
    'Dynamics'  : ['log2FC_Mfn1','log2FC_Mfn2','log2FC_Opa1',
                   'log2FC_FIS-1','log2FC_DRP-1'],
    'OxPhos'    : ['log2FC_COX','log2FC_CyBa','log2FC_CyC1',
                   'log2FC_ATP6','log2FC_NADHSub4','log2FC_NADHsub1']
}

print("\n=== PATHWAY-LEVEL Z-SCORE ===")
for path, cols in groups.items():
    mask = df[cols].notna().all(axis=1)
    Z = df.loc[mask, cols].apply(zscore, axis=0)
    df.loc[mask, f"{path}_zscore"] = Z.mean(axis=1)
    sub = df.dropna(subset=[f"{path}_zscore",'load','radiation'])
    stats = sub.groupby(['load','radiation'])[f"{path}_zscore"]\
            .agg(mean='mean', sem='sem')\
            .reset_index()
    print(f"\n-- {path} stats --\n{stats.to_string(index=False)}")
    model = smf.ols(f"{path}_zscore ~ C(load)*C(radiation)", data=sub).fit()
    print(f"\n-- (Wildman et al.) ANOVA --\n{anova_lm(model,
typ=3)[['sum_sq','df','F','PR(>F)']]}")

# 9) Pathway PC1 composites and ANOVA
print("\n=== PATHWAY-LEVEL PCA PC1 ===")
for path, cols in groups.items():
    mask = df[cols].notna().all(axis=1)
    X = df.loc[mask, cols].values
    if X.shape[0] < 4:
        print(f"Skipping PCA for (Wildman et al.) (too few samples)")
        continue
    pca = PCA(n_components=1)
    pc1 = pca.fit_transform(X).flatten()
    df.loc[mask, f"(Wildman et al.)_PC1"] = pc1
    ve = pca.explained_variance_ratio_[0]*100
    print(f"\n-- (Wildman et al.) PC1 var={ve:.1f}% --")
```

```
    sub = df.dropna(subset=[f"(Wildman et al.)_PC1",'load','radiation'])
    stats = sub.groupby(['load','radiation'])[f"(Wildman et al.)_PC1"]\
             .agg(mean='mean', sem='sem')\
             .reset_index()
    print(stats.to_string(index=False))
    model = smf.ols(f"(Wildman et al.)_PC1 ~ C(load)*C(radiation)", data=sub).fit()
    print(f"\n-- (Wildman et al.) PC1 ANOVA --\n{anova_lm(model,
typ=3)[['sum_sq','df','F','PR(>F)']]}")
```
