## Supplemental Table 1 for "Differential regulation of mitochondrial quality control in skeletal muscle by HZE radiation exposure and partial weightbearing in mice"

**Supplemental Table 1.** Results Statistical analysis of mRNA. A two-way ANOVA was conducted on all genes with main effects of Radiation (SHAM vs RAD) and Load (CC or 6/G. P values for the main effect and interactions are shown. ⁑ denotes data failed either Shapiro-Wilk test for normality or Levene's test for equal variance and therefore statistical analysis were conducted on power-transformed data using either Box-Cox or Yeo-Johnson. * denotes significant main effects.

| Genes | ME Radiation | ME Load | Interaction Load x Radiation |
|---|---|---|---|
| **Biogenesis** | | | |
| Nrf2 | 0.106 | 0.858 | 0.554 |
| PGC-1a | 0.035 * | 0.535 | 0.641 |
| Tfam | 0.051 * | 0.453 | 0.754 |
| Taco1 | 0.297 | 0.807 | 0.927 |
| Biogenesis Pathway | 0.051 * | 0.61 | 0.15 |
| **Autophagy** | | | |
| Beclin | 0.258 | 0.34 | 0.116 |
| LC3b | 0.033 * | 0.129 | 0.088 |
| p62 | 0.485 | 0.0669 | 0.341 |
| Gabarapl ⁑ | 0.473 | 0.689 | 0.263 |
| Park2 | 0.007 * | 0.856 | 0.98 |
| Bnip3 ⁑ | 0.972 | 0.239 | 0.052 * |
| Autophagy Pathway | 0.039* | 0.47 | 0.49 |
| **OXPHOS** | | | |
| ND1 ⁑ | 0.821 | 0.132 | 0.632 |
| ND4 ⁑ | 0.821 | 0.107 | 0.409 |
| CyBa | 0.499 | 0.904 | 0.682 |
| CyC1 | 0.556 | 0.731 | 0.085 |
| COX3 ⁑ | 0.262 | 0.799 | 0.125 |
| ATP6 ⁑ | 0.671 | 0.181 | 0.683 |
| OXPHOS Pathway | 0.110 | 0.248 | 0.098 |
| **Dynamics (Fusion/Fission)** | | | |
| Mfn1 ⁑ | 0.539 | 0.46 | 0.409 |
| Mfn2 ⁑ | 0.380 | 0.088 | 0.359 |
| Opa1 | 0.552 | 0.73 | 0.333 |
| Fis1 | 0.376 | 0.330 | 0.749 |
| Drp1 ⁑ | 0.780 | 0.568 | 0.459 |
| Dynamics Pathway | 0.662 | 0.270 | 0.311 |
